# Renal cancer cells acquire immune surface protein through trogocytosis and horizontal gene transfer

**DOI:** 10.1101/2024.08.07.607036

**Authors:** Haley Q. Marcarian, Anutr Sivakoses, Anika M. Arias, Olivia C. Ihedioha, Benjamin R. Lee, Maria C. Bishop, Alfred L.M. Bothwell

## Abstract

Trogocytosis is an underappreciated phenomenon that shapes the immune microenvironment surrounding many types of solid tumors. The consequences of membrane-bound proteins being deposited from a donor immune cell to a recipient cancer cell via trogocytosis are still unclear. Here, we report that human clear cell renal carcinoma tumors stably express the lymphoid markers CD45, CD56, CD14, and CD16. Flow cytometry performed on fresh kidney tumors revealed consistent CD45 expression on tumor cells, as well as varying levels of the other markers mentioned previously. These results were consistent with our immunofluorescent analysis, which also revealed colocalization of lymphoid markers with carbonic anhydrase 9 (CAIX), a standard kidney tumor marker. RNA analysis showed a significant upregulation of genes typically associated with immune cells in tumor cells following trogocytosis. Finally, we show evidence of chromosomal DNA being transferred from immune cells to tumor cells during trogocytosis. This horizontal gene transfer has transcriptional consequences in the recipient tumor cell, resulting in a fusion phenotype that expressed both immune and cancer specific proteins. This work demonstrates a novel mechanism by which tumor cell protein expression is altered through the acquisition of surface membrane fragments and genomic DNA from infiltrating lymphocytes. These results alter the way in which we understand tumor-immune cell interactions and may reveal new insights into the mechanisms by which tumors develop. Additionally, further studies into trogocytosis will help push the field towards the next generation of immunotherapies and biomarkers for treating renal cell carcinoma and other types of cancers.

**SIGNIFICANCE STATEMENT:** We have identified trogocytosis as a mechanism by which human clear cell renal carcinoma tumors acquire lymphocyte surface protein expression from tumor infiltrating immune cells. In addition to the transfer of membrane fragments, we have provided evidence to show that genomic DNA is transferred from a normal immune cell to a tumor cell during trogocytosis. This process alters the transcriptome of cancer cells such that they express significantly more mRNA for immune proteins such as the lymphocyte marker CD45 compared to tumor cells that have not undergone trogocytosis. This study provides an in-depth analysis of the interactions between cancer cells and tumor infiltrating lymphocytes, and how these interactions alter the development of human tumors.

## INTRODUCTION

Trogocytosis is the transfer of membrane fragments from a donor cell to an acceptor cell [1]. In mammalian cells, this process was first described on CD8^+^ T cells, in which the CD8^+^ T cells were shown to take fragments of the plasma membrane from certain antigen presenting cells (APCs) during the formation of the immunological synapse [2]. Since then, trogocytosis has been observed in many different types of immune cells, including CD4^+^ T cells [3], Natural Killer (NK) cells [4], and macrophages [5]. Previously, we have shown that trogocytosis plays an important role in the acquisition of immune regulatory molecules by colon cancer cells from infiltrating lymphocytes [6]. This resulted in increased immune regulator molecules such as CTLA4, PDL1, Tim3, VISTA, LAG3, CD38, CD80, CD86, MHC Class II, and PD-L1 being presented on the surface of trogocytic tumor cells compared to non-trogocytic tumor cells. Trogocytosis was also observed with numerous lymphocyte populations including macrophages, dendritic cells, NK cells, and monocytes[6]. Along with the transfer of surface proteins, genomic DNA PCR analysis also demonstrates the transfer of DNA from donor immune cells to mouse CRC cells, implying that tumor cells can display an altered transcriptome because of trogocytosis. Trogocytosis has also been implicated in other types of disease such as Hodgkin lymphoma and renal cell carcinoma[7, 8]. In mouse models of leukemia, trogocytosis promotes the transfer of target antigens from cancer cells to T cells, which induces T cell exhaustion and fratricide [9]. T cell exhaustion and fratricide emphasize the observation of relapse in patients treated with CAR T cell therapies. However, the extent to which trogocytosis occurs in tumors as well as the implications it has in shaping the tumor-immune microenvironment have not been fully elucidated.

Horizontal gene transfer (HGT) is a process that has been widely studied in the field of microbiology. HGT between different species of bacteria has been shown to grant survival advantages to the recipient microbe, such as resistance to antibiotics[10]. However, the concept of horizontal gene transfer between mammalian cells is largely uninvestigated. Recently, it has been shown that RPE1 cells, an immortalized non-transformed pigmented epithelium line can integrate a genomically encoded fluorescent reporter from a breast cancer cell line expressing that reporter when the two lines are cocultured[11]. Interestingly, physical contact was required between the two cell lines for HGT to occur. This indicates that HGT could occur concurrently with trogocytosis. To our knowledge, this is the first evidence of HGT between mammalian cells, however the mechanism behind such a transfer remains unknown. The consequences of this transfer, particularly in the context of cancer development and progression, remain uninvestigated.

In this study, we detected the presence of the immune markers CD14, CD16, CD56, and CD45 on tumor cells from renal cell carcinoma (RCC) patients. Previously, these markers have not been identified on RCC cells or other solid tumors. We suggest they were acquired through trogocytosis of infiltrating immune cells as these tumors develop. Other markers, including CD19 and CD11b were also tested and determined to not be significant markers of trogocytosis. These results were determined through quantification of immunofluorescent staining and flow cytometry. RNA analysis further showed trogocytosis between RCC cells and infiltrating lymphocytes resulted in an altered phenotype of tumor cells that expressed both tumor-specific and immune-specific genes. NanoString analysis of trogocytic tumor cells from human RCC tumors showed that trogocytic cells display rampant expression of immune cell-associated and tumor-associated genes. Finally, in vitro analysis of EdU and a genomically encoded GFP tag transfer from T cells to RCC cells showed that genomic DNA is transferred during the process of trogocytosis.

## RESULTS

### Identification and Quantification of Tumor Cells Presenting both Tumor and Lymphocyte Markers in Human RCC Samples

Macrophages have been recently implicated in abating the effects of immune checkpoint inhibitors (ICI) in treating RCC through the process of trogocytosis with cancer cells [8]. Given that RCC is an immunogenic cancer with high levels of immune infiltration, we hypothesized that other types of immune cells may play an important role in shaping the tumor microenvironment through trogocytosis. We obtained paraffin embedded slides from 21 human RCC tumors of varying stages to determine whether markers of trogocytosis could be detected. Positive staining for carbonic anhydrase 9 (CAIX) was used to define tumor cells as it is a well-characterized marker of hypoxia and is commonly upregulated in RCC cells [12].

To identify trogocytic tumor cells, slides were stained with CAIX and various immune markers of interest. Fluorescent imaging revealed significant populations of cells displaying both CAIX and CD14, CD16, CD56, or CD45RA (Figure 1A). Imaging of normal kidney tissue revealed no expression of these markers by healthy kidney epithelial cells (Supplemental Figure 1A). These results revealed that macrophages, monocytes, NK cells, and CD45RA^+^ T cells are likely the most active participants of trogocytosis with RCC cells. The RA isoform of CD45 is typically expressed on naïve CD4^+^ and CD8^+^ T cells, until it is converted into the RO form upon the formation of memory cells [13]. This implies that naïve T cells could be more trogocytic than mature memory T cells in RCC.

**Figure 1.**
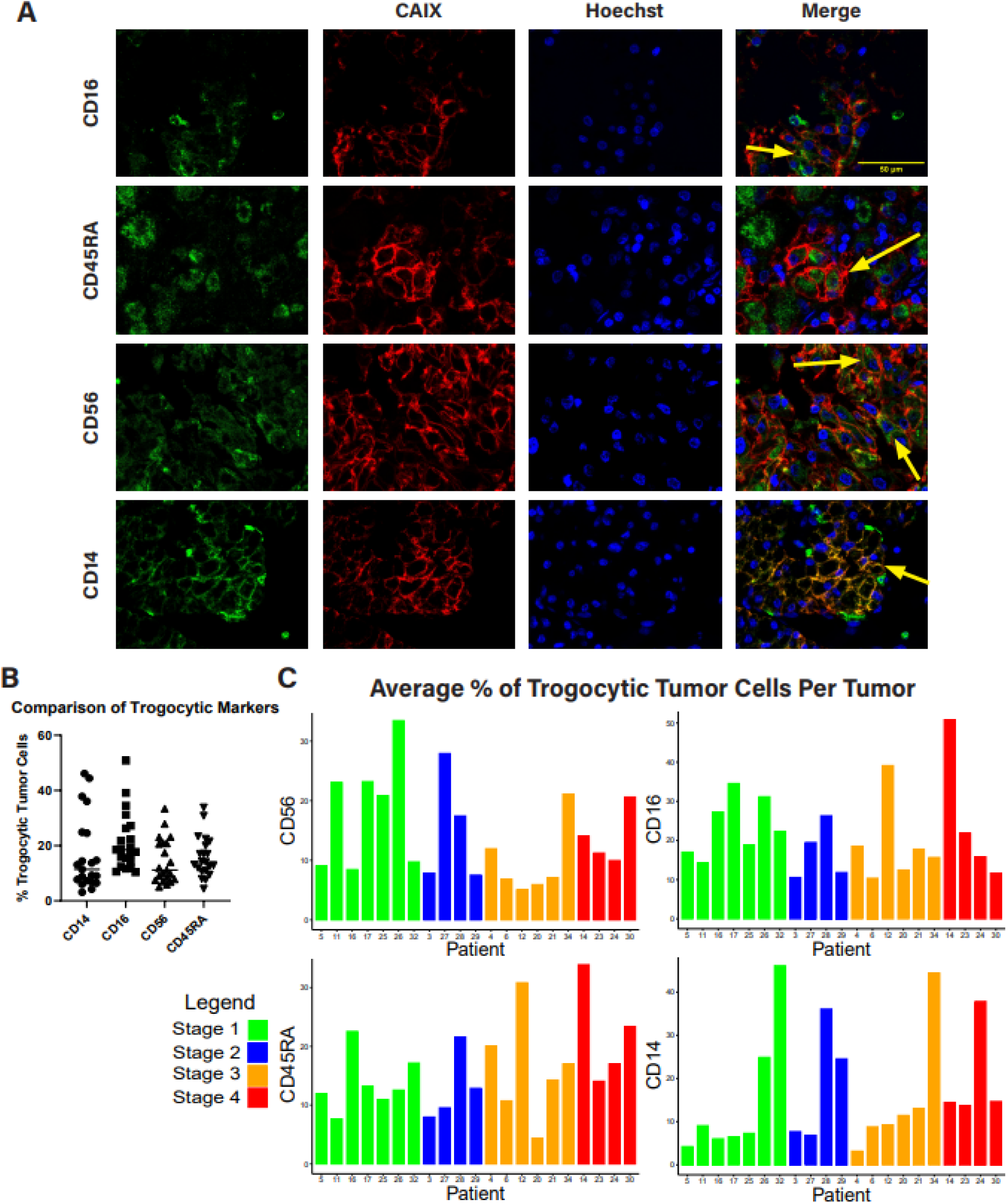
Trogocytic RCC tumor cells are detected and quantified using immunofluorescent staining. **(A)** Immunofluorescent microscopy of human RCC tumors. Trogocytic markers (CD14, CD56, CD45RA, CD16), CAIX, and Hoechst are displayed as green, red, and blue respectively. Arrows indicate trogocytic tumor cells. **(B)** Quantification of average percentage of trogocytic tumor cells for 21 human RCC tumors. **(C)** Quantification of average percentage of trogocytic tumor cells in 21 human RCC tumors organized by tumor stage. Methodology of quantification described in Methods and Materials.

To compare the percentage of tumor cells displaying signs of trogocytosis, we developed an image quantification program that records the percentage of individual cells’ surface area occupied by a trogocytic marker based on IF images (Supplemental Figure 1B). Cutoffs were established to define a tumor cell as trogo^+^ or trogo^-^ based on what estimated percentage of their surface area was positive for a given immune marker. Through this analysis, we found that of the 21 patient tumor images, all displayed significant signs of trogocytosis between immune cells and tumor cells (Figure 1 B,C). The average percentage of trogo^+^ RCC cells per tumor was between 15-20%, with some patient tumors as high as 50% trogo^+^ for certain markers.

### Characterization of Trogocytic RCC Tumors by Flow Cytometry

To demonstrate the transfer of CD45 transfer from a lymphocyte to a tumor cell in vitro we performed several coculture assays between RCC cell lines and Jurkat T cells or primary CD3^+^ T cells isolated from healthy donor PBMC. Cell lines were cocultured for 16 hours and then CD45 transfer was assessed via flow cytometry. Our results reveal that there was a significant increase in CD45 labeling on RCC cells that had been cocultured with primary T cells compared to RCC cells in monoculture (Figure 2A,B). We also observed a significant increase in CD45 expression by RCC cell in cocultures done with Jurkat T cells (Supplementary Figure 2A-C). This transfer was inhibited when a transwell barrier was placed between the two cell types in coculture, indicating that physical contact is necessary (Supplemental Figure 2D-E). We utilized size discrimination as well as CAIX expression to ensure no T cells were included when determining whether a CD45^+^ cell was truly a trogocytic cancer cell (Supplementary Figure 2F-G).

**Figure 2.**
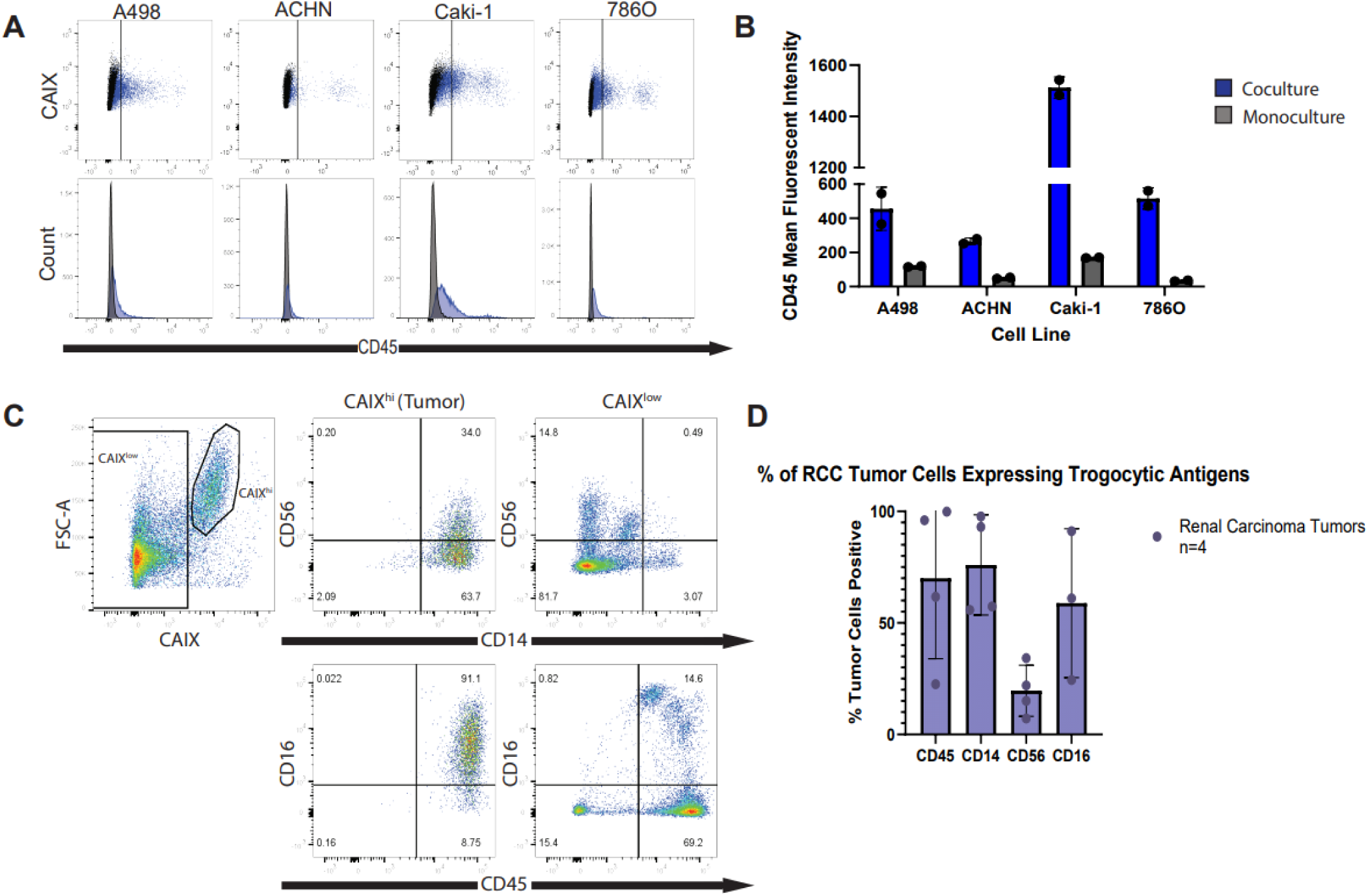
RCC cells acquire expression of immune surface proteins through contact with lymphocytes. **(A)** Flow cytometry analysis of CD45 expression by RCC cell lines post coculture with primary human T cells. RCC cells were cocultured with T cells for 16 hours prior to analysis. Data shows an increase in CD45 expression relative to RCC cells cultured in isolation (monoculture). **(B)** Mean fluorescent intensity of CD45 expression by cocultured RCC cells relative to monoculture. **(C)** Representative flow cytometry analysis of fresh human ccRCC tumors. Tumor cells were identified as CAIX^hi^ and lymphocytes and other cells present in the tumor microenvironment were identified as CAIX^low^. Data shows expression of four immune cell surface proteins (CD14, CD16, CD56, CD45) by ccRCC and non-ccRCC cells isolated from this tumor. **(D)** Percentage of tumor cells from 4 human ccRCC tumors that expressed CD14, CD16, CD56, and CD45 based on flow cytometry analysis. Tumor cells were identified via CAIX^hi^ labeling as represented in Figure 2C.

Next, we further characterized fresh human RCC tumors for markers of trogocytosis via flow cytometry. The IF analysis previously described has certain limitations that make distinguishing trogocytic staining from background signal difficult. Flow cytometry was chosen to provide a more accurate depiction of the trogocytic properties of RCC tumor cells. Fresh tumor samples were acquired from the University of Arizona biobanking facility and immediately dissociated and stained for markers of interest, namely CAIX, CD45, CD56, CD16 and CD14. Samples were evaluated within 12 hours of receiving them to ensure maximum viability of the cells.

Tumor cells were discriminated from infiltrating lymphocytes and non-tumor tissue gating on CAIX^hi^ cells. Size discrimination was also used to determine that the CAIX^hi^ population was independent from lymphocyte populations present in the tumor (Figure 2C). After establishing the tumor cells as distinct from all other cell types present in the tissue, we further characterized both the tumor and lymphocyte populations using a panel of trogocytic markers (CD14, CD16, CD56, and CD45). Analysis showed significant levels of expression of all four markers by fresh human ccRCC tumor cells (Figure 2C,D, Supplementary figures 3A,4).

CD45 is an established marker for all nucleated cells of hematopoietic lineage[14]. CD45 labeling revealed widespread expression on the surface of the CAIX^hi^ (tumor) population (Figure 2C,D). Our data suggest that the majority of tumor cells in RCC tumors have expression of immune markers, and that this property is detectable by flow cytometry.

Given the high percentage of tumor cells exhibiting trogocytic markers, we then decided to focus on characterizing cells that express multiple immune proteins. Our data revealed that the vast majority of CAIX^hi^ cells were positive for multiple immune cell markers (Figure 2D). Therefore, it is possible that RCC tumor cells undergo multiple trogocytic events that allow them to “steal” the ability to express these markers from multiple types of immune cells. The exact identity of these immune cells is not yet fully elucidated and is an important field of study for future investigations.

### RCC Tumor cells Express RNA for Immune Surface Proteins

Given the abundance of immune antigens being expressed by RCC tumor cells, we decided to identify the source of this protein expression. Upwards of 98% of tumor cells in human RCC tumors were positive for one or more trogocytic protein when characterized by flow cytometry (Figure 2C). The current definition of trogocytosis describes the transfer of membrane fragments from one cell to another, however it seemed unlikely that simple membrane transfer from lymphocytes to tumor cells could account for the relatively high quantities of trogocytic protein present on the surface of RCC cells. Therefore, we sought to identify other possible sources of this protein expression by tumor cells. More specifically, we attempted to determine whether tumor cells were capable of self-sufficient transcription of immune cell-specific proteins after trogocytosis occurred. To investigate this, we cocultured Jurkat T cells with four kidney carcinoma cell lines. We performed fluorescence activated cell sorting (FACS) to isolate RCC cells from the coculture prior to harvesting RNA to ensure there was no contamination of T cells in the cells being analyzed. Using qRT-PCR analysis, we identified a significant induction of CD45 gene expression in RCC cells post coculture (Figure 3C).

**Figure 3.**
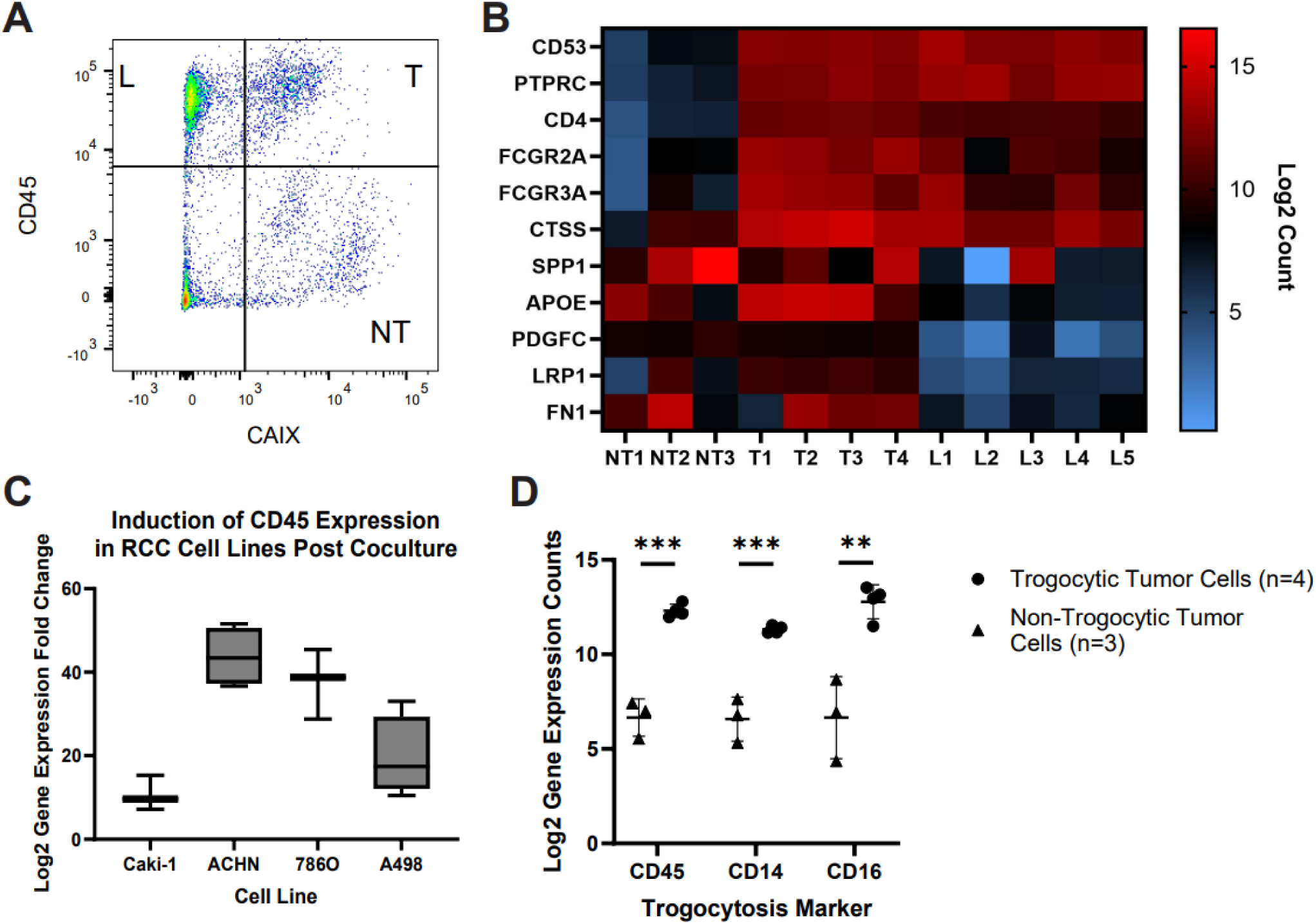
RNA expression analysis of immune cell-specific genes by trogocytic RCC cells in human ccRCC tumors. **(A)** Representative gating strategy used to identify lymphocyte (L), trogocytic tumor (T), and non-trogocytic tumor (NT) cell populations in human ccRCC tumors when sorting cells via FACS. L cells were identified as CD45high CAIX^low^, trogocytic tumor cells were identified as CD45^high^ CAIX^high^, and non-trogocytic tumor cells were identified as CD45^low^ CAIX^high^. **(B)** NanoString® gene expression analysis for human ccRCC tumor. Populations were sorted from fresh human ccRCC tumors via FACS according to the gating strategy described in Figure 3A. Data shows Log2 RNA expression counts for select immune and cancer specific genes. **(C)** qRT-PCR analysis of the change in CD45 expression in RCC cell lines post coculture with Jurkat T cells. Cells were sorted via FACS prior to analysis to isolate RCC cells. Graph depicts Log2 CD45 gene expression fold change in cocultured cells relative to monoculture. (Mean Ct values (n=4): Caki-1 coculture 28.297, monoculture 31.174, ACHN coculture 29.131, monoculture 33.575, 786O coculture 28.501, monoculture 36.759, A498 coculture 29.071, monoculture 33.385). (D) Comparison of Log2 gene expression counts of trogocytosis markers between trogocytic and non-trogocytic tumor cell populations isolated from human RCC tumors. Data was analyzed using multiple T tests (p value < 0.01 = **, p value < 0.001 = ***).

Next, we isolated trogocytic (T, CAIX^hi^CD45^hi^), non-trogocytic (NT, CAIX^hi^CD45^low^), and lymphocyte (L, CAIX^low^CD45^hi^) populations from human RCC tumor samples via FACS sterile sorting (Figure 3A). We then ran an nCounter® PanCancer Immune Profiling analysis on the sorted cells using Nanostring technology to identify key transcriptional differences between T and NT cells. Our analysis showed that trogocytic tumor cells exhibit a “fusion” phenotype, defined as expressing both tumor and immune-specific genes (Figure 3B). There was significantly higher expression of the trogocytic markers CD45, CD14, and CD16 in trogocytic tumor cell populations compared to non-trogocytic tumor cell populations (Figure 3D). We also identified new potential trogocytic markers such as CD4 and CD53 on the trogocytic tumor samples (Figure 3B, Supplementary Table 1).

### RCC Cell Lines Acquire Genomic DNA from Primary T Cells

In order to determine the mechanism by which RCC tumor cells gained immune protein expression, we investigated the possibility of horizontal gene transfer (HGT) occurring during trogocytosis. To do this, we generated a primary T cell line expressing GFP-tagged histone H2B as a secondary method of tracking DNA transfer. GFP-H2B T cells were cocultured overnight with Caki-1 cells and then analyzed via flow cytometry. Analysis showed that there was a small population of GFP-positive Caki-1 cells that was not present in monocultured Caki-1 cells (Figure 4A). We observed a significant increase in the mean fluorescence intensity of GFP in cocultured Caki-1 cells compared to monocultures of Caki-1 cells alone (Figure 4B).

**Figure 4.**
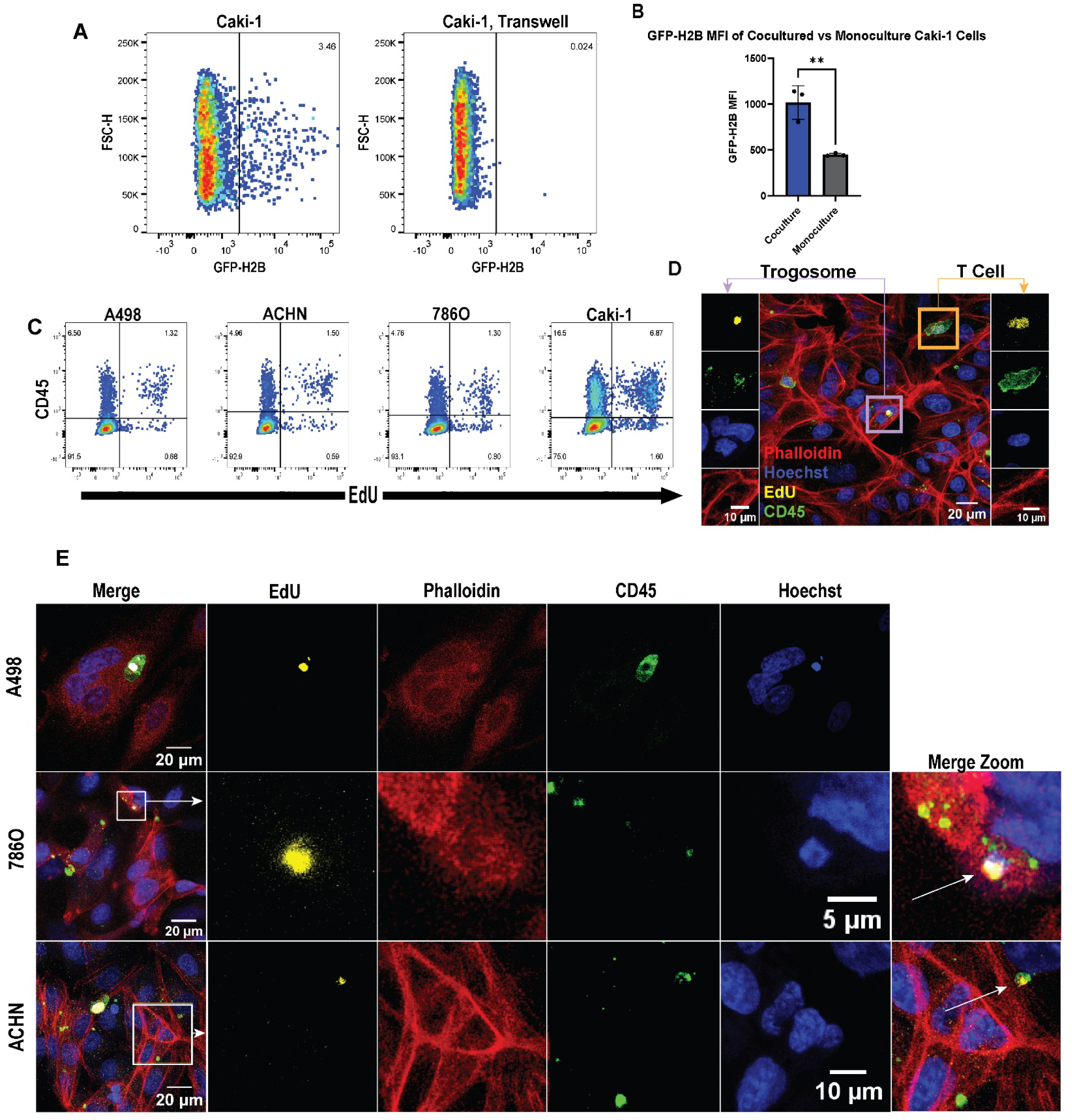
DNA is transferred from primary T cells to RCC cell lines. **(A)** Flow cytometry results of Caki-1 cells that were cocultured with GFP-histone H2B primary T cells. Data shows the percentage of GFP-H2B^+^ Caki-1 cells in coculture relative to monocultured Caki-1 cells. **(B)** Mean fluorescent intensity of GFP-H2B expression of cocultured Caki-1 cells compared to monoculture (p value = 0.0058, unpaired T test was used to analyze the data). **(C)** Flow cytometry analysis of RCC cell lines cocultured with EdU-labeled primary human T cells. Plots show the percentage of RCC cells that are EdU^+^ and CD45^+^ post 16-hour coculture. **(D)** Image taken from a 10:1 coculture of EdU-labeled primary human T cells with ACHN cells. Left panels depict an enhanced view of an EdU^+^ CD45^+^ trogosome within the intracellular space of an ACHN cell. Right panels depict an enhanced view of an EdU^+^ CD45^+^ T cell for comparison purposes. **(E)** Representative images of trogosomes in three RCC cell lines (top to bottom, A498, 786O, ACHN). Boxes in the merged images of middle and bottom panels indicate the region being enhanced in the split color panels to the right of the merged image. Arrows in the merged zoom panels indicate the location of a trogosome. All images taken on a Zeiss 710 laser scanning confocal, 40X (NA 1.30).

Additionally, we isolated primary human T cells from a healthy donor and incubated them with EdU, a thymidine analog that is readily incorporated into the DNA of dividing cells[15]. EdU-labeled T cells were then cocultured overnight with RCC cell lines (ACHN, A498, Caki-1, and 786O cells). Following coculture, the cells were then analyzed via flow cytometry to determine whether we could detect a transfer of EdU-labeled DNA from the T cells to the RCC cell lines. Results showed that between 1-8% of cocultured RCC cells contained EdU labeled DNA (Figure 4C). As contamination from EdU labeled T cells could result in false positive results, we used size discrimination in the flow cytometry analyses to distinguish cancer cells from T cells (Supplementary figure 5A,B). In contrast, monocultured RCC cell lines and cocultures done with a transwell control separating the T cells from the RCC cells did not show any evidence of DNA transfer (Supplementary Figure 5C).

Finally, we performed super-resolution structured illumination microscopy to gain insight into the physical properties of the transferred material. We discovered CD45^+^ intracellular bodies present in RCC cells post coculture with primary human T cells (Figure 4D,E). Additionally, some of these “trogosomes” contained EdU labeled DNA (Figure 4D,E). Interestingly, we observed instances in which there was a lack of colocalization between EdU and Hoechst DNA labeling (Figure 4E, ACHN). This is likely due to the covalent attachment of bulky chemical groups to DNA as a result of EdU labeling that can block Hoechst from binding, particularly in A-T rich regions of DNA[15]. The presence of these trogosomes could be indicative of a mechanism by which lymphocyte DNA is transferred through CD45^+^ lymphocyte membrane bubbles into the intracellular space of RCC cells.

## DISCUSSION

In this study, we showed that RCC tumor cells express significant levels of lymphoid proteins, such as CD45RA, CD14, CD16, and CD56. We have demonstrated that this expression is obtained through trogocytosis and subsequent horizontal gene transfer occurring between a tumor cell and lymphocyte. Importantly, this phenomenon is widespread, with over 90% of fresh RCC tumor cells showing high levels of trogocytic protein expression when analyzed via flow cytometry. Additionally, RNA extraction and analysis showed that tumor cells exhibit an altered transcriptome after trogocytosis occurs. Namely, we observed an increase in RNA expression for genes that are typically only expressed by cells of hematopoietic lineage. Finally, we demonstrated that genomic DNA can be transferred during trogocytosis from a lymphocyte to a tumor cell resulting in an altered tumor cell phenotype. Taken together, this data demonstrates that RCC tumor cells acquire the ability to express large quantities of hematopoietic surface antigens following a trogocytic event with infiltrating lymphocytes.

Using immunofluorescent analysis of RCC slides, we developed a computer algorithm to quantify the percentage of tumor cells that express proteins found on lymphocytes. This study revealed consistent expression of the markers CD14, CD16, CD56, and CD45RA in a cohort of 21 human patient tumors when analyzed via immunofluorescence microscopy (IF). Despite limitations in the sensitivity of this method, analyzing 21 tumors originating from different stages of disease revealed no significant difference in trogocytic antigen expression at any stage of tumor development. Additionally, of the four trogocytic antigens we analyzed, there were no significant changes in the specific type of antigen being expressed at any stage. These data imply that trogocytosis occurs at an early stage during tumor development, and that the trogocytic phenotype of RCC tumor cells are maintained as disease progresses. Currently, it is unknown whether this is due to repeated trogocytic interactions with infiltrating lymphocytes, acquisition of lymphocytic DNA that promotes expression of these antigens, or some combination of the two.

Analysis of fresh tumors by flow cytometry corroborated the results we obtained via IF. Flow cytometry revealed much higher levels of trogocytic marker expression than previously anticipated, with over 90% of tumor cells in all RCC tumors tested being positive for at least one lymphocytic marker. To our knowledge, this is the first time these lymphocytic proteins have been identified on any solid tumor and could shift the paradigm of how we understand a tumor’s interactions with its microenvironment.

Our investigation into horizontal gene transfer between RCC cells and T cells has shown that it is possible for tumor cells to acquire DNA from lymphocytes that they encounter. We have shown that the transfer of both EdU-labeled DNA and GFP-tagged histones is observed after coculturing tumor cells and immune cells *in vitro*. Currently, it is unknown whether this transfer confers greater survival capabilities or a more aggressive phenotype to tumor cells that have undergone horizontal gene transfer. Additional studies are needed to fully characterize this process and its role in disease progression. Our data also suggests that membrane transfer can occur independently of horizontal gene transfer, implying that there are multiple mechanisms that regulate these processes.

Trogocytosis has been shown to promote the acquisition of the immune check point receptor PD-1 by natural killer cells in mouse models of leukemia, resulting in suppressed NK cells antitumor immunity [16]. Additionally, colorectal cancer cells have been shown to obtain other immune regulatory molecules such as CTLA4 through trogocytosis with infiltrating lymphocytes [6]. These results suggest that trogocytosis is a mechanism by which tumors evade elimination by the immune system, either by acquiring regulatory molecules themselves, or by indirectly benefiting from certain immune cells negatively regulating themselves. In the context of RCC, it is unknown how tumor cells use the acquisition of immune cell surface protein and DNA to promote their survival. However, given the frequent expression of these markers on RCC tumors from all stages of disease, we believe trogocytosis confers a survival advantage to tumor cells. More research is needed to determine whether this survival advantage is selected for by the tumor to evade the immune system, or some other unknown mechanism.

It is possible that trogocytosis and the effect it has on the phenotype of tumor cells may be exploited to improve cancer therapies. The expression of lymphoid antigens on cancer cells may constitute effective targets of antibody drug conjugates (ADCs). Further profiling of the markers that are transferred during this process may reveal unique protein expression signatures by trogocytic tumors that are not otherwise found in healthy cells. In addition to their use as new drug targets, these markers can also potentially be used as novel biomarkers to help better inform treatment strategies for clear cell renal carcinoma patients. Further research in the field of trogocytosis is critical for developing the next generation of immunotherapies.

## ACKNOWLEDGEMENTS

We acknowledge use of the University of Nebraska Medical Center - UNMC Advanced Microscopy Core Facility, RRID:SCR_022467, P20 GM103427 (NIGMS, NE-INEBRE), P30 GM106397 (NIGMS, NCS), P20GM130447 (NIGMS, CoNDA), P30 CA036727 (NCI, Buffett Cancer Center), S10RR02730 (NIH), S10OD030486 (NIH), Nebraska Research Initiative, UNMC Vice Chancellor for Research Office. Research reported in this publication was supported by the National Cancer Institute of the National Institutes of Health under award number P30 CA023074. The UNMC Flow Cytometry Research Facility is administrated through the Office of the Vice Chancellor for Research and supported by state funds from the Nebraska Research Initiative (NRI) and The Fred and Pamela Buffett Cancer Center’s National Cancer Institute Cancer Support Grant (P30 CA036727). Major instrumentation has been provided by the Office of the Vice Chancellor for Research, The University of Nebraska Foundation, the Nebraska Banker’s Fund, and by the NIH-NCRR Shared Instrument Program. Special thanks to J.C. Fitch for assistance with flow cytometry analysis. Studies were supported by funding from the National Cancer Institute (T32CA009213-44) to A. Sivakoses and 7R01AI137060-06 awarded to A.L.M. Bothwell.

## USE OF HUMAN MATERIALS

All human materials were acquired through the University of Arizona Tissue Acquisition and Cellular/Molecular Analysis Shared Resource. Patient materials were obtained and processed in accordance with University of Arizona tissue acquisition policy.

## DATA AVAILABILITY

Source code for immunofluorescent analysis available for use at bit.Ly/TrogoTracker.

## MATERIALS AND METHODS

### Immunofluorescent staining assays

The following primary antibodies were used in immunofluorescent experiments: anti-human CAIX (R&D Biosystems, AF2188), CD45RA (Sino Biological Inc., 102580-T08), CD56 (ProteinTech®, 14255-1-AP), CD68 (Sino Biological Inc., 11192-T26), CD16 (ProteinTech®, 16559-1-AP), CD14 (R&D Biosystems, BAF383). Paraffin-embedded tissue slides were obtained from Tissue Acquisition and Cellular/Molecular Analysis (TACMASR) core facility at the University of Arizona. Slides were deparaffinized for 9min in Xylenes. They were then washed in decreasing concentrations of ethanol solutions at 1min intervals for 5min. The slides were then washed in MilliQH_2_O for 10 minutes. Then, antigen retrieval was performed using boiling Antigen Unmasking Solution (Vector Laboratories, H-3300) for 10min and left to cool for an additional 20min. Slides were blocked with 20% donkey serum/1X DPBS-0.1% Tween-20 for 30min. Primary antibody solutions were prepared according to their recommended dilutions. Slides were incubated with primary antibodies overnight at 4°C and washed with 1X DPBS-0.1% Tween-20 for 5min. Samples were quenched using Invitrogen ReadyProbes™ Tissue Autofluorescence Quenching Kit for 3-5min. Secondary anti-donkey antibodies from Invitrogen were applied and incubated for 1 hour at room temperature in the dark. Samples were quenched an additional time and stained with Hoescht nuclear stain for 10min. Coverslips were mounted on the stained slides using Fluoromount-G™ (Invitrogen, 00-4958-02). Slides were imaged using the Echo Inc. Revolution fluorescent microscope.

### Flow cytometry assays

The following conjugated antibodies were used in flow cytometric experiments: CD45 (clone 2D1, BioLegend, 368532), CD16 (clone CB16, Invitrogen, 17-0168-42), CD14 (clone 61D3, Invitrogen, 414-0149-42), CD45RO (clone UCHL1, Invitrogen, 12-0457-42), CD45RA (clone HI100, Invitrogen, 17-0458-42), CD44 (clone IM7, Invitrogen, 404-0441-82), CAIX (R&D Biosystems, FAB2188G), CD56(clone CMSSB, Invitrogen, 12-0567-42), and CD68 (clone REA886, Miltenyi Biotec, 130-114-463). Fresh tumor tissue was acquired through the Tissue Acquisition and Cellular/Molecular Analysis (TACMASR) core facility at the University of Arizona. Tumor samples were finely minced into 1mm^2^ sections and washed with DPBS. They were then suspended in a solution of 1mg/ml Collagenase Type 4 (Worthington, LS0004186) and incubated at 37°C for 30min, being shaken every 3min. The reaction was stopped using DMEM media with 5% fetal bovine serum. Cell suspensions were then filtered through a 40µm mesh filter and resuspended in 100µl of DPBS. Cell viability was stained with Ghost Dye™ Violet 450 (Tonbo Biosciences, 13-0863-T100). Human BD Fc Block™ (BD Pharmingen™, 564220) was used at a concentration of 5µl/10^6^ cells for 10min at room temperature. Fluorescent antibody staining was done at the recommended concentrations of each antibody for 30min at room temperature in the dark. Cell suspensions were then washed using DPBS and resuspended in 1X RBC lyse/fix solution (eBioscience™ 00-5333-57) for 40min. Samples were then washed in DPBS and analyzed using the BD FACSCanto™ II system. Data was analyzed using FlowJo v10.8.1.

### Immunofluorescent image quantification

Images used for analysis were taken at 40X (NA 1.40) on an Echo Revolution LED microscope. Images were exported in separate channels into ImageJ (FIJI) for quantification. Nuclei were identified using the StarDist2D plugin [17], then a Voronoi algorithm was applied to estimate the boundaries of individual cells. The tumor marker channel was used to subtract all non-tumor cells from the Voronoi cell map. The resulting regions of interest (ROIs) were then saved to be used as a map for all tumor cells in each image. The channel containing the trogocytic marker of interest was then run through a background subtraction algorithm and the resulting signal was normalized using the CLAHE histogram normalization plugin. The channel was then converted to a binary image and the ROIs previously described were applied. The total surface area of each cell was measured to give a two-dimensional representation of the cellular surface area being occupied by a trogocytic marker. Cells with trogocytic markers occupying over 55% and less than 90% of their total 2D surface area were considered trogocytic. These cutoffs were determined to exclude autofluorescence, other sources of background signal, and infiltrating immune cells from quantification. We believe this method of quantification is quite conservative and may exclude low expressing trogocytic tumor cells, implying that the true percentage of tumor cells that are trogocytic may be much higher than this analysis estimates. This could account for the difference between the IF quantification and flow cytometry results.

### Cell Culture

Human primary T cells were obtained from the Elutriation Core Facility at the University of Nebraska Medical Center. T cell populations were isolated using the Miltenyi Biotec Pan T Cell Isolation Kit (130-096-535). T cells were cultured in X-VIVO™ 15 Serum-Free Hematopoietic Cell Medium (Lonza, 02-053Q) supplemented with 20ng/mL recombinant human IL-2 (R&D, BT-002-010) once a week and 7ul/mL ImmunoCult™ Human CD3/CD28/CD2 T Cell Activator (Stem Cell Technologies, 10970) every other week. RCC cell lines (Caki-1, ACHN, A498, 786O) were obtained from Dr. George Sutphin’s laboratory at the University of Arizona. RCC cell lines were cultured based on the recommendations made by ATCC.

### Lentiviral Transduction of Primary T Cells

GFP-H2B lentiviral vector was obtained from Dr. Ghassan Mouneimne at the University of Arizona. Lentivirus was generated in 239T cells, and the viral supernatant was harvested for primary T cell infection. T cells were spun for 30 minutes at 3000 rpm with the virus. Lentivirus was removed the following day after incubation and GFP^+^ T cells were isolated using FACS selection 2 weeks after transduction.

### Coculture Assays

RCC lines (ACHN, A498, Caki-1, 786-O) were plated a minimum of 4 hours prior to coculture. T cells were added to the culture at a 1:1 ratio and incubated overnight. Cocultures were harvested for experimental purposes 16 hours post addition of the T cells.

### Confocal Microscopy of EdU Transfer

Images were taken on a Zeiss 710 Laser Scanning Confocal microscope at 40X (NA = 1.30). Images shown are max intensity projections of 60 Z-stacked images. FIJI smoothing algorithm was applied to phalloidin imaging for the purpose of minimizing the quality loss of actin labeling as a result of the EdU staining process.

