## Supplemental Figure 1 for "Renal cancer cells acquire immune surface protein through trogocytosis and horizontal gene transfer"

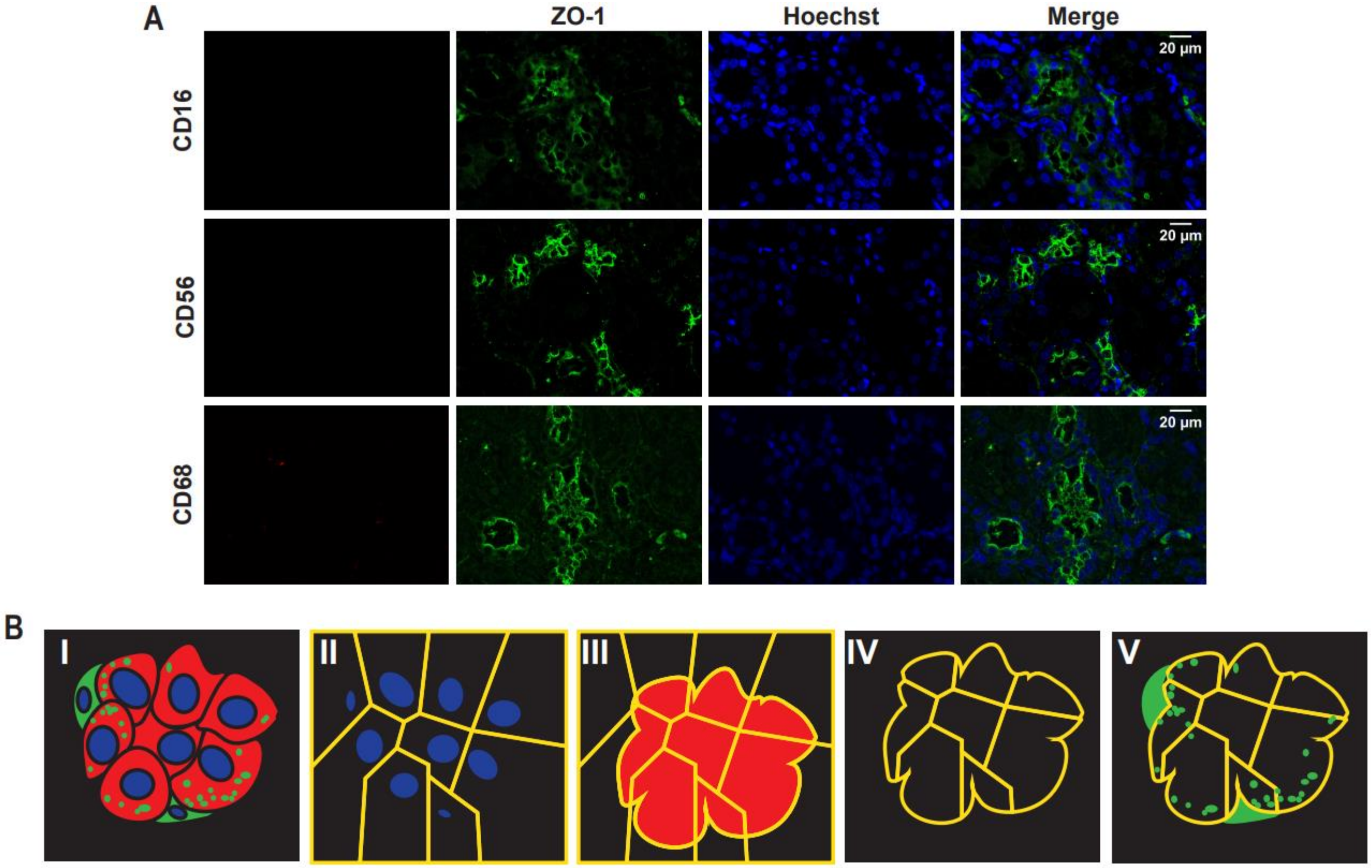


**Supplemental Figure 1. (A)** Immunofluorescent staining of normal kidney tissue. Trogocytic markers shown in the left panel, ZO-1 staining highlights normal kidney epithelial cells. **(B)** Representative illustration of the algorithm used to quantify immunofluorescent kidney slides. **(B,I)** Cancer cells represented in red (CAIX), lymphocytes represented in green (CD45). Trogocytic tumor cells represented by expression of both CAIX and CD45. **(B,II)** Detection of nuclei through hoechst staining, Voronoi diagram is applied based on the location of nuclei and segments cells. **(B,III and IV)** CAIX staining is applied to the previous voronoi diagram and used to exclude non-CAIX$^+$ cells such as lymphocytes and non-tumor tissue. **(B,V)** Final voronoi diagram is applied to CD45 labeling to approximate the boundaries of cells and determine the expression levels of CD45 on tumor cells only.
