## Supplemental Figure 2 for "Renal cancer cells acquire immune surface protein through trogocytosis and horizontal gene transfer"

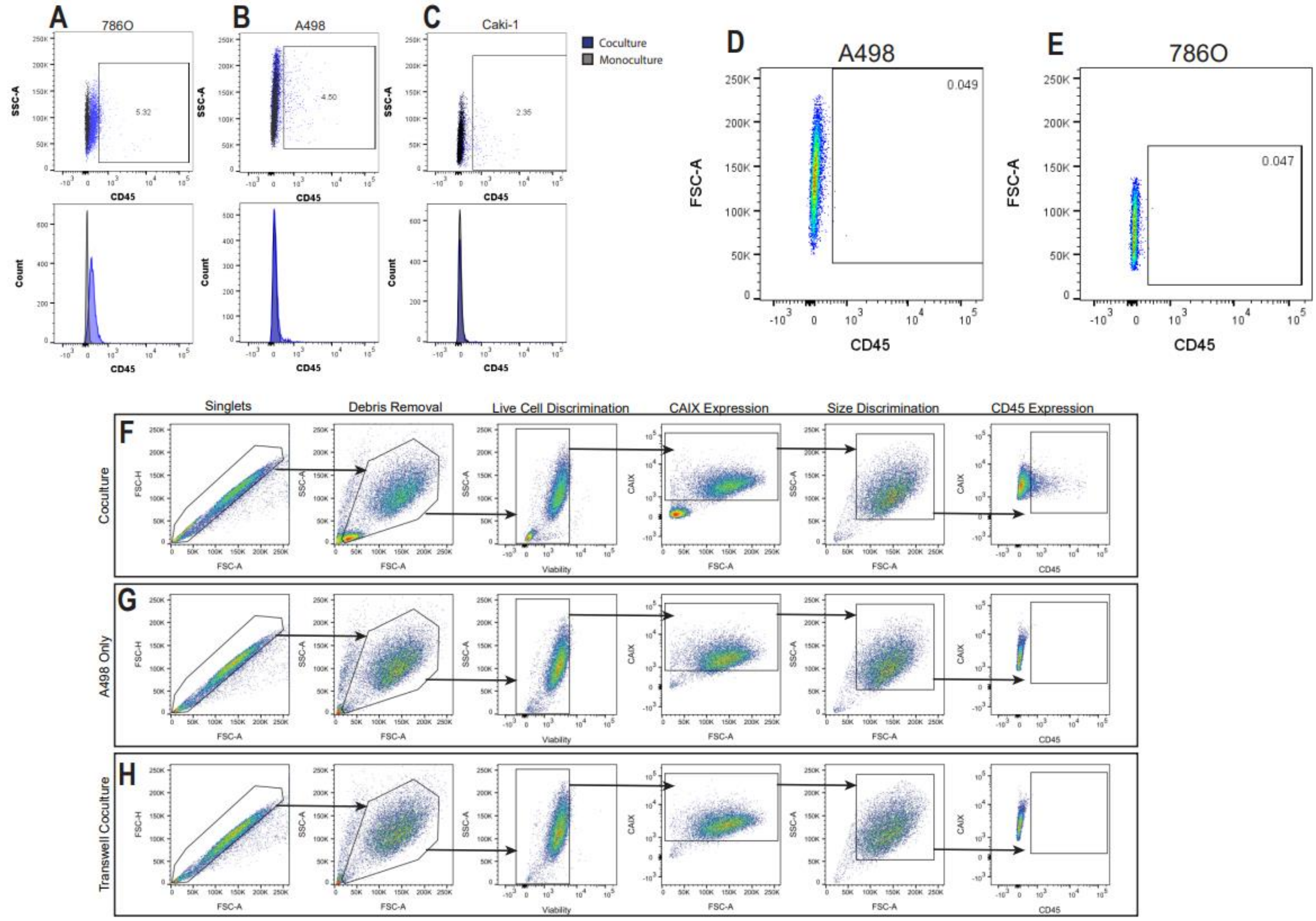


**Supplemental Figure 2. (A-C)** Flow cytometry analysis of RCC cell lines that were cocultured with Jurkat T cells to asses the transfer of CD45. (Top) gating indicates the percentage of RCC cells that are CD45$^{+}$ relative to monoculture controls. **(D,E)** Flow cytometry analysis of A498 and 786O cocultures with primary human T cells that were separated by a transwell barrier. Gating indicated the percentage of RCC cells that were positive for CD45 post-over night coculture. **(F-H)** Flow cytometry gating strategy used to remove T cell for analysis of CD45 positive cancer cells. Example depicts A498 cells cocultured with primary T cells **(F)**, A498 cells only **(G)**, and A498 cells separated from primary T cells using a transwell barrier **(H)**.
