## Supplemental Figure 3 for "Renal cancer cells acquire immune surface protein through trogocytosis and horizontal gene transfer"

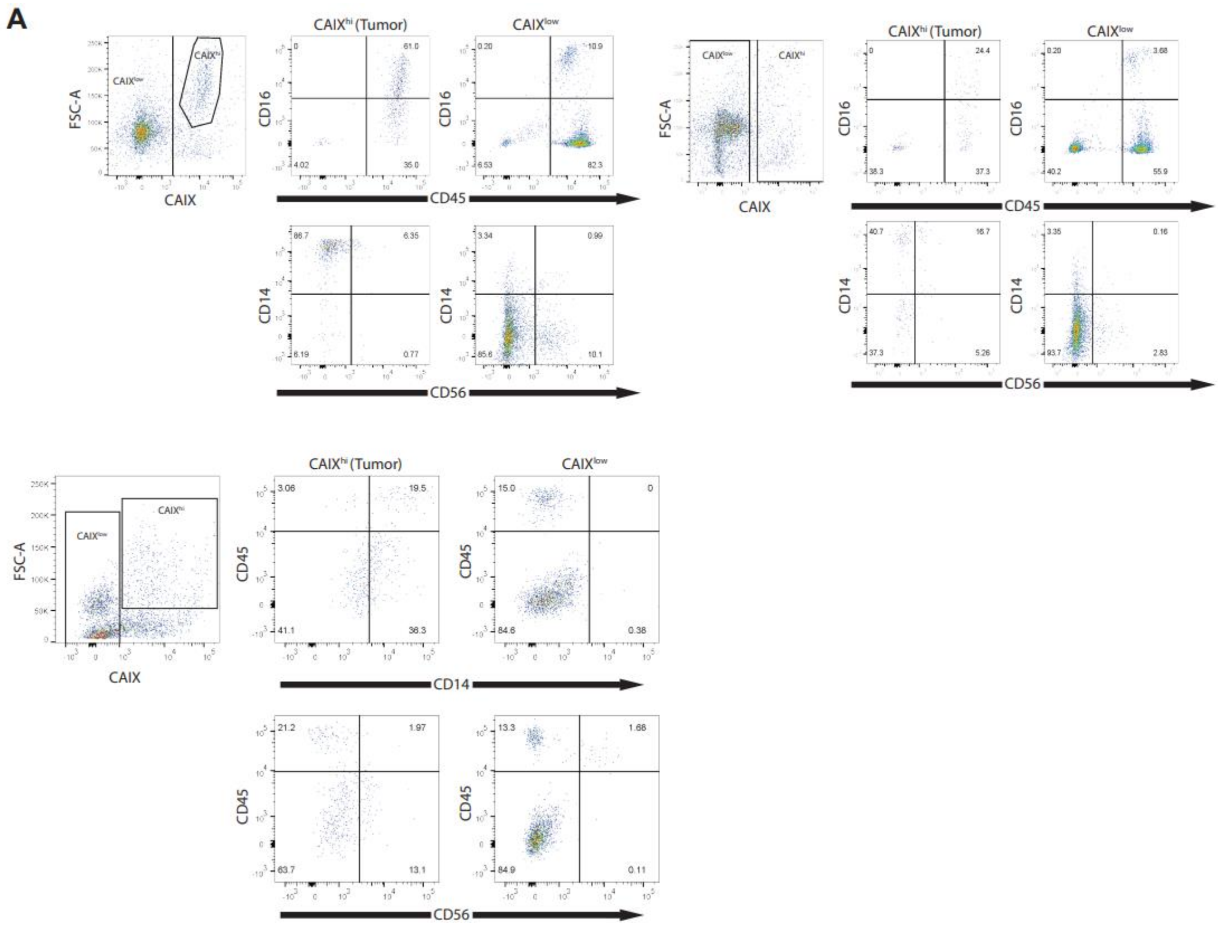


**Supplementary Figure 3.** Flow cytometry results of three additional fresh human RCC tumors. Left plots indicate the gating strategy used to isolate CAIX$^+$ tumor cells and CAIX$^{low}$ lymphocytes. The right plots for each tumor indicate the percentage of cells in these two populations that are positive for the indicated immune cell proteins.
