## Supplemental Figure 4 for "Renal cancer cells acquire immune surface protein through trogocytosis and horizontal gene transfer"

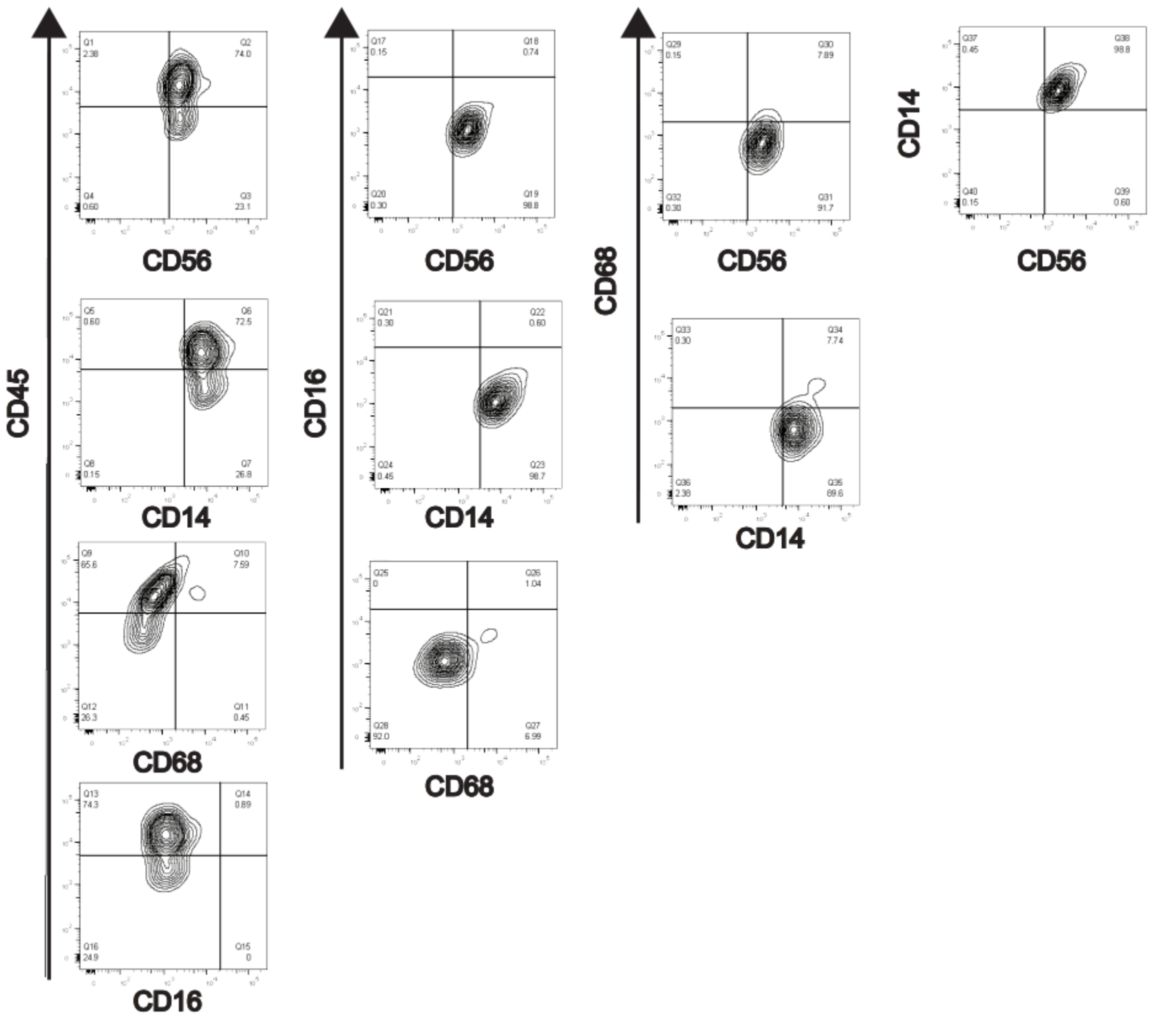


**Supplemental Figure 4.** Multiparametric representation of an additional fresh kidney tumor flow cytometry data based on expression of the trogocytosis markers CD45, CD14, CD16, CD56, and CD68. Prior to analysis, tumor cells were isolated based on CAIX$^+$ expression, and size discrimination.
