## Supplemental Table 1 for "Renal cancer cells acquire immune surface protein through trogocytosis and horizontal gene transfer"

| Trogocytic Tumor vs Non-trogocytic Tumor | | Trogocytic Tumor vs Lymphocytes | |
|---|---|---|---|
| **Gene ID** | **Log2 Ratio** | **Gene ID** | **Log2 Ratio** |
| CD48 | 7.83 | C3 | 6.82 |
| CCL3L1 | 7.81 | TREM2 | 6.73 |
| LY86 | 7.68 | TLR7 | 6.44 |
| C3AR1 | 7.62 | CD163 | 6.39 |
| HCK | 7.48 | APOE | 6.36 |
| LILRB3 | 7.47 | CXCL12 | 6.35 |
| CLEC7A | 7.37 | A2M | 6.22 |
| CSF1R | 7.33 | C1QB | 6.00 |
| CD86 | 7.08 | C1QA | 5.85 |
| CCR5 | 7.07 | MSR1 | 5.71 |
| TREM2 | 6.91 | F13A1 | 5.47 |
| TLR7 | 6.87 | SIGLEC1 | 5.32 |
| CD33 | 6.85 | CASP10 | 5.17 |
| LAIR2 | 6.81 | CSF1R | 5.04 |
| CD84 | 6.60 | PDGFC | 5.01 |
| CLEC5A | 6.60 | TLR5 | 4.87 |
| S100A12 | 6.58 | C2 | 4.85 |
| CSF3R | 6.57 | CD14 | 4.71 |
| FCER1G | 6.57 | LRP1 | 4.55 |
| CCR2 | 6.53 | CXCL10 | 4.45 |

**Supplementary Table 1.** Top 20 differentially expressed genes detected in the Nanostring analysis of fresh ccRCC tumors. (Left) Results show a comparison of the most differentially upregulated genes in trogocytic tumor cells relative to non-trogocytic tumor cells based on the PanCancer Immune Profiling Nanostring® panel. (Right) Top 20 differentially upregulated genes in trogocytic tumor cells relative to tumor infiltrating lymphocytes. Results are based on mean gene expression counts of 3 non-trogocytic tumor cell populations, 4 trogocytic tumor cell populations, and 5 tumor infiltrating lymphocyte populations isolated from human ccRCC tumors.
