## Supplemental Figure 5 for "Renal cancer cells acquire immune surface protein through trogocytosis and horizontal gene transfer"

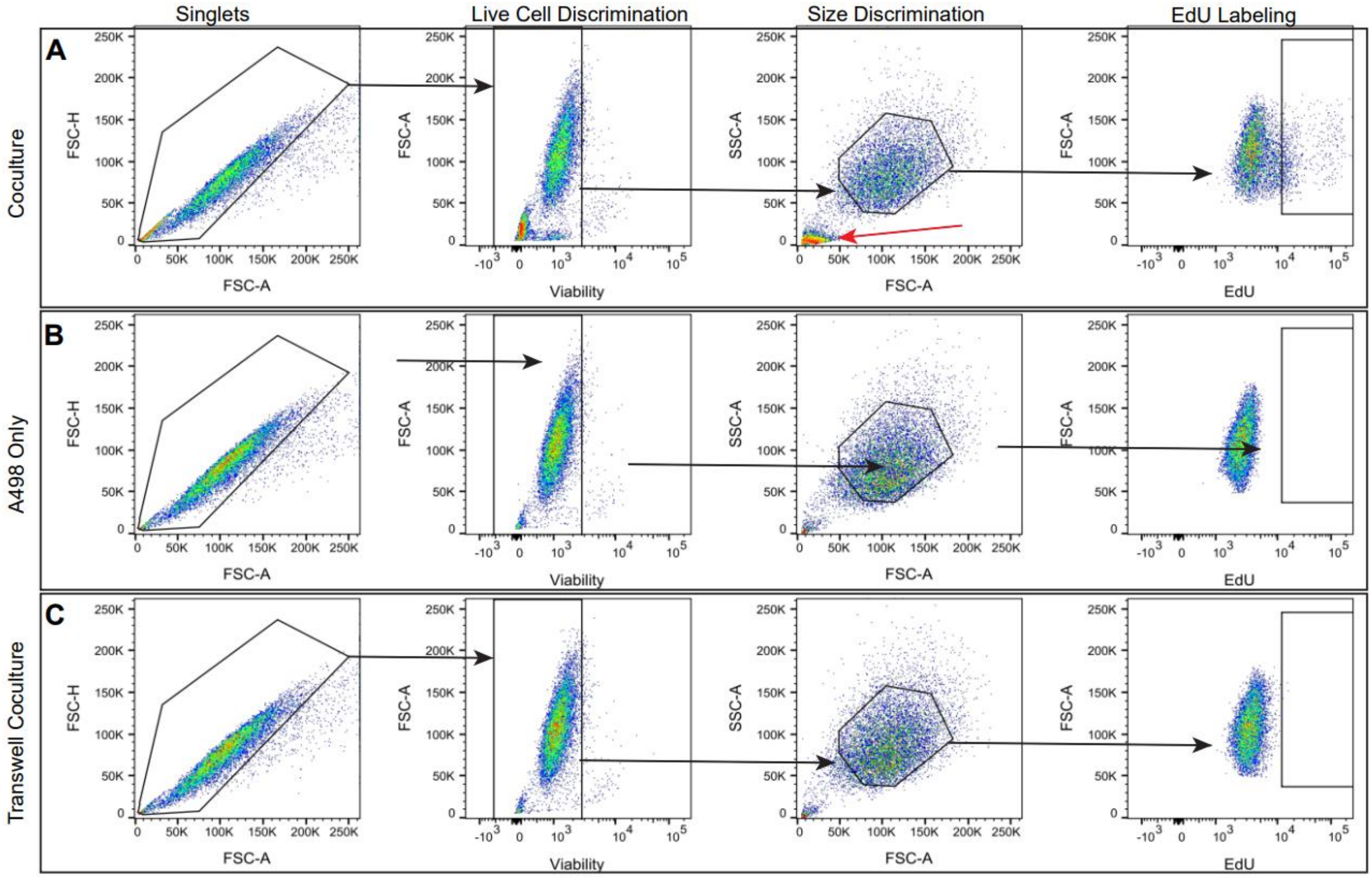


**Supplemental Figure 5.** Comprehensive gating strategy used to identify EdU positive RCC cells (A498 depicted). **(A)** Single cells in coculture suspension were identified using FSC-H FSC-A discrimination. Live cells were determined using UV Ghost Dye 450 as a marker for viability. SSC-A and FSC-A were used for identifying T cells and RCC cells, subsequently isolating RCC cells only for analysis. Red arrow indicates the position of T cells. FSC-A and EdU plot depicts the gate used to identify EdU positive cancer cells. **(B)** Gating strategy applied to a monoculture of only A498 cells. **(C)** Gating strategy applied to an A498-T cell coculture that was separated by a transwell barrier. T cells were removed with the barrier prior to flow cytometry analysis.
